# Multiple stages of evolutionary change in anthrax toxin receptor expression in humans

**DOI:** 10.1101/2020.07.29.227660

**Authors:** Lauren A. Choate, Gilad Barshad, Pierce W. McMahon, Iskander Said, Edward J. Rice, Paul R. Munn, James J. Lewis, Charles G. Danko

## Abstract

The advent of animal husbandry and hunting increased human exposure to zoonotic pathogens. To understand how a zoonotic disease influenced human evolution, we studied changes in human expression of anthrax toxin receptor 2 (*ANTXR2*), which encodes a cell surface protein necessary for *Bacillus anthracis* virulence toxins to cause anthrax disease. In immune cells, *ANTXR2* was 8-fold down-regulated in all available human samples compared to non-human primates, indicating regulatory changes early in the evolution of modern humans. We also observed multiple genetic signatures consistent with recent positive selection driving a European-specific decrease in *ANTXR2* expression in several non-immune tissues affected by anthrax toxins. Our observations fit a model in which humans adapted to anthrax disease following early ecological changes associated with hunting and scavenging, as well as a second period of adaptation after the rise of modern agriculture.

---

Despite abundant evidence that infectious diseases have driven human adaptation (*1*–*8*), we understand relatively little about how exposure to new pathogens can shape human host genetics. Here we study the evolution of human host gene expression to *Bacillus anthracis* infection, a bacterium that causes anthrax disease (*9, 10*), as a model for how repeated intermittent exposure to an infectious disease can drive recurrent adaptive alteration. *B*. *anthracis* primarily afflicts ruminants during the course of its natural life-cycle, and has driven selective pressures in cattle and sheep (*11, 12*). While anthrax disease is believed to rarely affect most primate species, with exceptions (*13, 14*), anthrax disease has been a notable source of mortality in humans (*9, 10*). Genetic diversity of extant *Bacillus* strains suggests that anthrax-causing *Bacillus* pathogens evolved in sub-Saharan Africa (*15, 16*), likely well before human radiation around the globe (*17*). *B*. *anthracis* then radiated from Africa into the middle east and Europe approximately 3-6ka, possibly following the spread of Neolithic agricultural practices (*16*). *B*. *anthracis* strains endemic to Europe later spread around the globe through trade and colonization (*16*). Thus, although anthrax disease has been associated with the rise of animal husbandry (*18*– *20*), the putative African origin of anthrax disease-associated *Bacillus* species may have provided opportunities for early humans to encounter *B*. *anthracis*, or the ancestral *B*. *cereus*, while hunting or scavenging ruminate game species well before the advent of modern farming. This led us to hypothesize that humans may have adapted to anthrax disease during several periods of human evolution—first to early ecological changes associated with hunting and scavenging, followed by a second period of adaptation after the rise of agriculture.

To identify evolutionary differences associated with anthrax susceptibility in humans, we studied changes in CD4+ T-cells between humans and non-human primates. We confirmed that in T-cells anthrax toxins inhibit cellular activation using an ELISA for the T-cell activation marker IL2 (**Fig. S1**), consistent with studies indicating that anthrax toxins inhibit immune responses that could clear the pathogen (*21*–*23*). We then studied transcriptional changes between human and non-human primate CD4+ T-cells using PRO-seq (Pol II loading) and RNA-seq (mature mRNA). We focused our analysis on the *ANTXR2* gene, which encodes a ubiquitously expressed transmembrane receptor that aids in the cellular entry of toxins secreted by the *B*. *anthracis* bacterium (*24*–*26*) (**Fig. 1A**). Consistent with our hypothesis, *ANTXR2* was 8-fold downregulated in human CD4+ T-cells compared to non-human primates at the level of both Pol II loading and mRNA (**Fig. 1B-C; Fig. S2A-B**). Expanding our analysis of RNA-seq data from CD4+ T-cells to 85 humans confirmed the loss of *ANTXR2* expression in all of the available human data (*27*) (**Fig. 1D**). Moreover, analysis of RNA-seq from peripheral blood mononuclear cells supported decreased *ANTXR2* expression in Batwa (a hunter gatherer population historically in the Great Lakes region of Africa) and Bakiga (agriculturalists from neighboring Rwanda and Uganda), compared to rhesus macaque (*28, 29*) (**Fig. S3**). Collectively, analysis of *ANTXR2* expression data was consistent with our hypothesis of ancient evolutionary changes affecting *ANTXR2* expression during the early divergence of humans from other primates.

**Figure 1.**
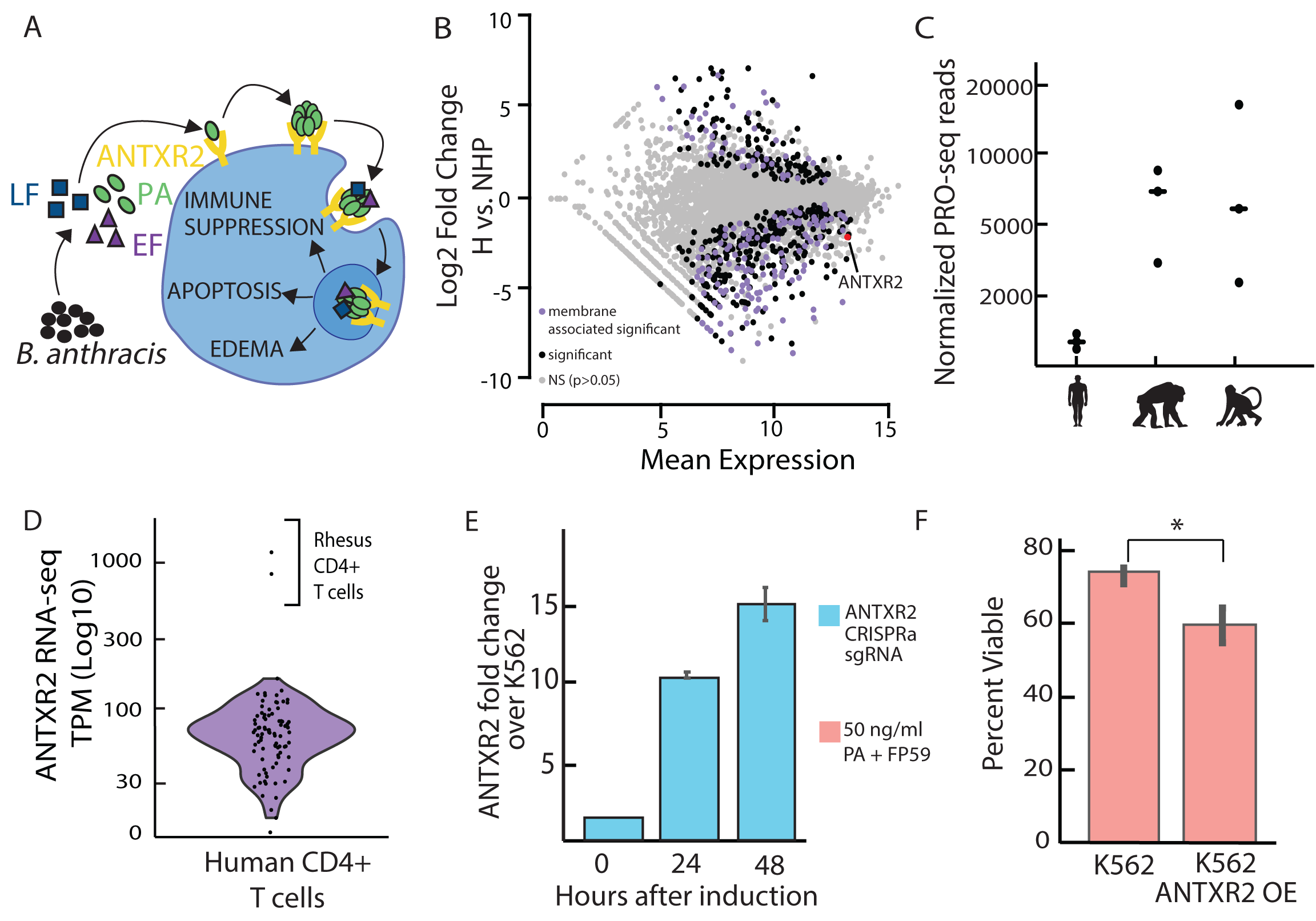
Decreased *ANTXR2* in humans can affect anthrax toxin sensitivity in blood cells. **A)** B. anthracis toxins lethal factor (LF) and edema factor (EF) cause apoptosis, edema, and immune suppression in target host cells. **B)** Differentially transcribed protein-coding genes between humans and non-human primates in CD4+ T cells based on PRO-seq data. The GO term ‘integral component of the membrane’ is enriched in differentially transcribed genes. **C)** CD4+ T cell PRO-seq data shows that *ANTXR2* is transcribed 8-fold lower in humans than chimpanzee and rhesus macaque. **D)** Comparison of *ANTXR2* RNA expression levels among a large set of humans (n=85) to rhesus macaque. Human variation in expression does not overlap with rhesus macaque expression. **E)** CRISPRa induction in K562 cells results in increased ANTXR2 expression as measured using qRT-PCR. F) CRISPRa K562 cells that overexpress *ANTXR2* are less viable after an anthrax toxin challenge as measured by the alamar blue viability stain.

To confirm that *ANTXR2* expression changes of the magnitude observed between human and non-human primates can affect sensitivity to anthrax toxins, we overexpressed *ANTXR2* using CRISPR activation (CRISPRa) (*30*). CRISPRa increased *ANTXR2* mRNA levels by 10-fold relative to an empty vector control 24 hours after transfection, a change that was similar in magnitude to the difference observed between humans and non-human primates (**Fig. 1E**). *ANTXR2* overexpressing cells had a significantly lower viability following treatment with recombinant anthrax toxins: protective antigen (PA) and FP59, a potent analog of lethal factor (LF) (*31*) (**Fig. 1F**). Thus a 10-fold increase in human *ANTXR2* expression, similar to changes found in non-human primates, increased the effect of anthrax toxins on cellular phenotypes.

We asked whether cis-regulatory changes could account for transcriptional divergence in *ANTXR2* between species. We used enhancer RNA transcription detected using PRO-seq data to identify candidate cis-regulatory elements (CREs) near *ANTXR2* in CD4+ T-cells of each primate species (*32*). dREG annotated 14 candidate CREs near *ANTXR2*, including four in introns of *ANTXR2*, a broad promoter, six CREs situated within the gene desert that separated *ANTXR2* and *PRDM8*, and three CREs within the region surrounding the *PRDM8* transcription unit (**Fig. 2A**). Notably, a comparison with DNase-I hypersensitivity and H3K27ac ChIP-seq data in humans did not identify additional candidate CREs in this locus (**Fig. S4**). Testing each putative CRE for differential transcription, a hallmark of regulatory activity at enhancers (*33*), revealed a strong bias for decreased transcription in the human lineage across the *ANTXR2* locus (median 2.3-fold lower transcription in human; *p* = 1e-3; Wilcoxon rank sum test; **Fig. S5**). At least three CREs were differentially transcribed on the human lineage using a conservative test for differential transcription (median CRE 16-fold lower in human; FDR corrected p-value < 0.05). Notably, the majority of CREs in the gene desert or near either the *ANTXR2* or *PRDM8* promoter were found in orthologous positions in both non-human primate species. Since the majority of genome-wide CRE activity changed between chimpanzee and rhesus macaque (*34*), the deep conservation of gene desert CREs may indicate an important biological function. The significant degree of cis-regulatory divergence between human and non-human primates at multiple annotated CREs thus led us to speculate that cis-acting loci, rather than trans-acting factors, underlie much of the divergence in *ANTXR2* expression.

**Figure 2.**
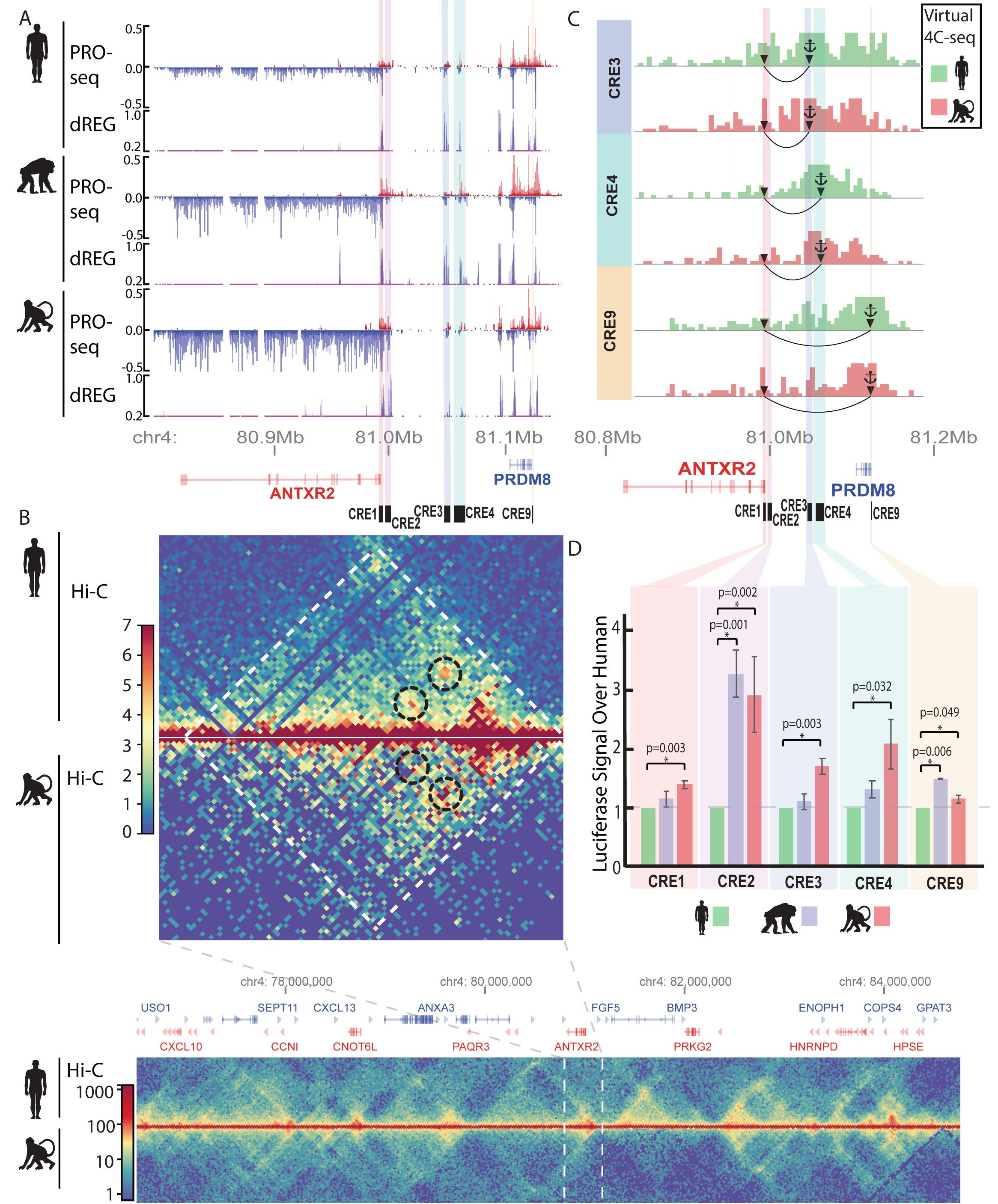
Changes in *ANTXR2* cis-regulatory element activity and chromatin structure. **A)** Genome browser shot of CD4+ T cell PRO-seq (normalized by reads per million) in human, chimpanzee, and rhesus macaque and regulatory elements and regulatory elements predicted by dREG. **B)** Hi-C in CD4+ T cells from human and rhesus macaque. *ANTXR2* is located in the same topological domain (TAD) as the upstream gene *PRDM8*. TADs are marked with white dotted lines. Focal contacts with the *ANTXR2* promoter are circled with black dotted lines. **C)** Virtual 4C-seq plots for distal regulatory elements tested in the luciferase assay show contacts to the *ANTXR2* promoter. **D)** Luciferase assay performed in Jurkat cells to test the activity of regulatory elements in human, chimpanzee, and rhesus macaque shows an increase in activity for either chimpanzee or rhesus macaque compared to humans for CRE1, CRE2, CRE3, CRE4, and CRE9.

While species-specific variation in *ANTXR2*-associated CREs is suggestive of CRE-mediated transcriptional evolution, we aimed to explicitly link distal CREs to the *ANTXR2* gene. To verify that annotated CREs interact with the *ANTXR2* promoter, we performed *in situ* Hi-C (*35*) in CD4+ T-cells from human and rhesus macaque and tested for enrichment of contacts in the *ANTXR2* locus. Our Hi-C data revealed that *ANTXR2, PRDM8*, and all candidate CREs were found within the same topological associated domain (TAD), a structure reported to insulate distal enhancers from affecting expression (*36, 37*) (**Fig. 2B**). Moreover, Hi-C data revealed focal contacts between the *ANTXR2* promoter and candidate CREs in the gene desert upstream (**Fig. 2B**). The main focal contacts observed in human Hi-C data had a higher contact frequency with the *ANTXR2* promoter in rhesus macaque, potentially reflecting species-specific differences in *ANTXR2* transcription (**Fig. 2C; Fig. S6**). Virtual 4C-seq analysis confirmed that several of the distal candidate CREs had increased contact frequency with the *ANTXR2* promoter relative to flanking regions (**Fig. 2C; Fig. S7**). Additionally, we identified an *ANTXR2* expression quantitative trait locus (eQTL) that overlapped CREs in the gene desert and near *PRDM8* using RNA-seq data from human CD4+ T-cells (*27*) (**Fig. S8**). Taken together, these findings indicate that many of the evolutionarily conserved CREs throughout the upstream gene desert interact with the *ANTXR2* promoter and affect *ANTXR2* expression in CD4+ T-cells.

Multiple CREs changed Pol II loading in human, which raised the possibility that multiple CREs contributed to changes in *ANTXR2* expression. To determine whether changes in the transcriptional activity at multiple CREs reflect independent DNA sequence changes, we used a luciferase assay to measure the activity of CREs using DNA sequences from human, chimpanzee, and rhesus macaque. We focused on 9 CREs in the upstream gene desert, the *ANTXR2* promoter, and in the *PRDM8* transcription unit, most of which were active in orthologous positions in multiple species, providing evidence of evolutionary constraint indicative of a biological function. We introduced nine CREs into Jurkat human leukemic CD4+ T-cells, a model trans-environment which recapitulates the pattern of transcription in the *ANTXR2* locus observed in primary T-cells (**Fig. S9**). Five of the 9 candidate CREs showed higher luciferase activity in chimpanzee or rhesus macaque than in human (*p* < 0.05, t-test; **Fig. 2D; Fig. S10**). Previous studies have noted multiple, compensatory changes in the activity of sequences within a CREs (*38*–*41*). Consistent with this idea, breaking CREs into separate regions in some cases uncovered compensatory changes within distinct regions of the CREs, for instance within the complex *ANTXR2* promoter (**Fig. S11**). Nevertheless, decreased overall activity in five of nine CREs, with no CREs increased in human, is unlikely to happen by chance (*p* = 0.03, binomial test; *p* = 0.01, multinomial test assuming 36% of CREs change (*41*)). Thus, our results may best fit an adaptive model in which human ancestors had an ancient adaptive change for reduced *ANTXR2* expression that is now fixed in humans.

In addition to changes compared with other primates, *ANTXR2* expression varied by one order of magnitude within humans, suggesting that a number of genetic variants may still exist within human populations associated with anthrax susceptibility. Most extant *B*. *anthracis* strains outside of Africa show genetic similarity to those in Europe, leading to a model in which *B*. *anthracis* that was endemic to Europe during the Neolithic era spread throughout the globe by trade and colonization during the past several thousand years (*16, 42*–*44*). Indeed, previous reports have noted lower anthrax toxin sensitivity in Europeans (*45*). To determine how selection has influenced modern human populations at the *ANTXR2* locus, we computed the composite-likelihood-ratio (CLR) of a selective sweep across chromosome 4 in four human populations (Europeans [CEU], East Asians [CHB and JPT], and Africans [YRI]) using SweepFinder2 (*46*). Consistent with the hypothesis that anthrax disease in Europeans led to a relatively recent population specific adaptive response, we found a candidate selective sweep in the gene desert upstream of the *ANTXR2* locus (**Fig. 3A**). The selective sweep had a higher CLR in European (CEU) than 98% of other loci on chromosome 4 (**Fig. S12**). Moreover, the CLR was substantially higher in Europeans compared to 1000 Genomes populations representative of East Asian (CHB, JPT) or African (YRI) ancestry (**Fig. 3A**). Surprisingly, this sweep had minimal overlap with CREs significantly associated with decreased *ANTXR2* expression between humans and non-human primates, and suggests a complex evolutionary response driven by recurrent selection around *ANTXR2*.

**Figure 3.**
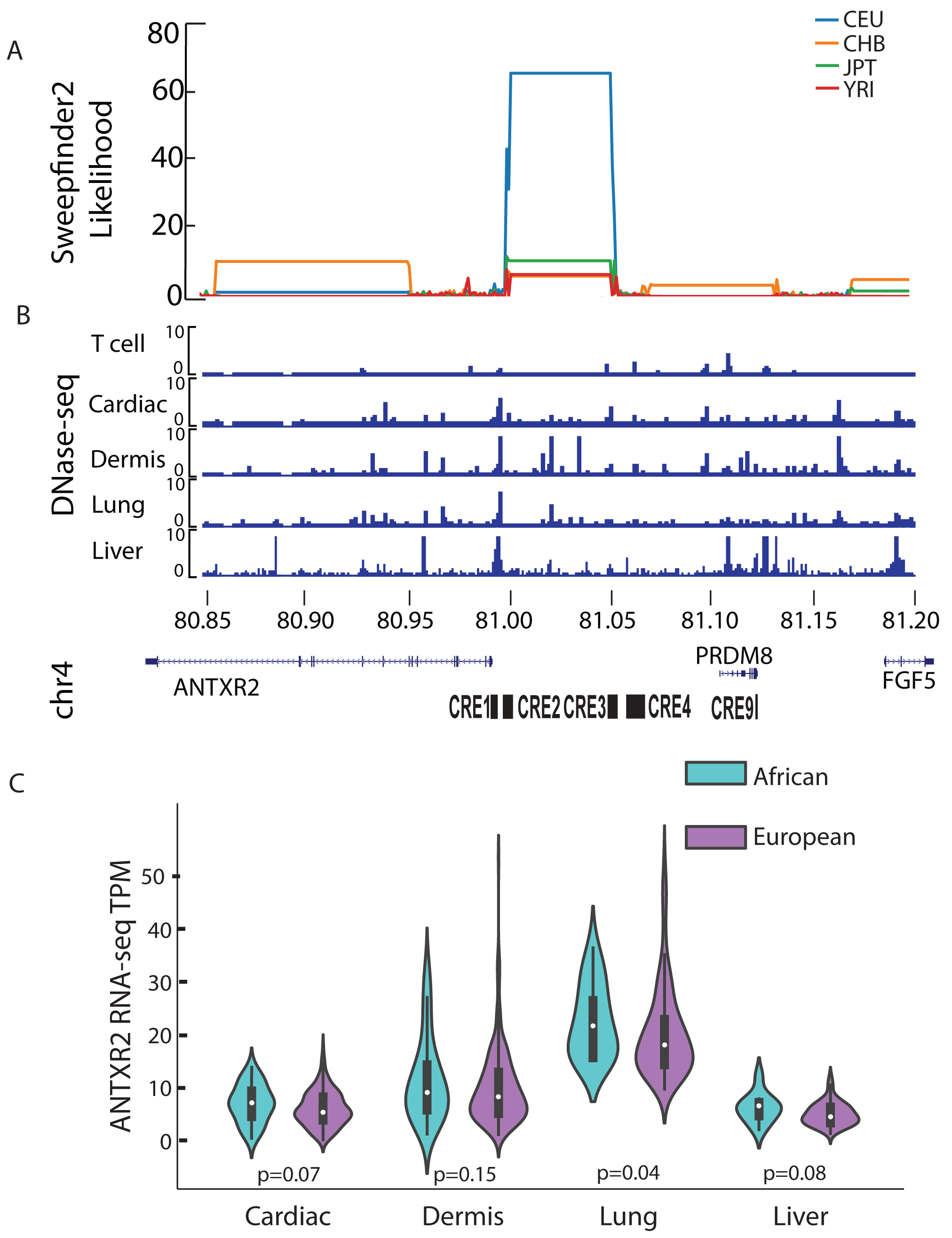
Hard selective sweep affects ANTXR2 expression in multiple tissues. **A)** Sweepfinder2 shows a predicted selective sweep in the CEU population upstream of *ANTXR2*. **B)** ENCODE DNase-I-seq data for T cells, cardiac cells, dermis, lung, and liver show differential regulatory landscapes within the predicted selective sweep in CEU. **C)** GTEX data for cardiac, dermis, lung, and liver show increased *ANTXR2* expression in African American individuals compared to European individuals.

The discovery of a selective sweep at the *ANTXR2* locus that excluded loci associated with decreased *ANTXR2* expression in CD4+ T-cells led us to speculate that a recent hard sweep may have occurred in response to anthrax-related selection pressures on additional organs. Anthrax disease is a systemic disorder affecting multiple organs in different manners. The main effects of anthrax disease are apoptosis (by anthrax lethal toxin), edema (by anthrax edema toxin), and a suppressed immune response (*26, 47*). The main targets of anthrax lethal toxin are cardiomyocytes and smooth muscle cells, whereas the main targets of anthrax edema toxin are the dermis, hepatocytes, and lungs (*48, 49*). Anthrax disease progresses with attacks on multiple organ systems, but the cause of host death is often by the induction of apoptosis in the heart (*50, 51*). We therefore examined human DNase-I hypersensitivity data in 42 tissues from the Epigenome Roadmap project (*52*). We found substantial numbers of DNase-I hypersensitive sites overlapping the interval of high CLR values in heart, lung, liver, and dermis (**Fig. 3B, Fig. S13**). Of these, multiple putative CREs were not found in blood DNase-I samples, supporting the hypothesis that the recent selective sweep affected tissues susceptible to apoptosis or edema.

Evidence of decreased expression between populations in non-blood tissues would be consistent with our prediction of recurrent selection on multiple disease-affected organs. To test whether *ANTXR2* expression varies among populations at anthrax disease targeted tissues, we divided GTEx RNA-seq data into European (n = 209, 306, 135, 60 for heart, skin, lung, and liver, respectively) and African (n = 28, 43, 17, 10 for heart, skin, lung, and liver) ancestry based on the mitochondrial haplotype (*53*). Although we were underpowered in most tissues due to the small number of individuals of African ancestry, *ANTXR2* expression was reduced in all four tissue types in Europeans relative to African samples and significantly reduced in lung (Mann-Whitney U test *p* = 0.07, 0.15, 0.04, 0.08 in heart, skin, lung, and liver; **Fig. 3C**). Combining p-values across tissues using Fisher’s method also supported decreased *ANTXR2* expression in Europeans (*p* = 0.0085, Fisher’s method). While additional study will be needed, we speculate that Europeans have, on average, decreased expression of *ANTXR2* in several tissues that are central to anthrax disease pathogenesis via recent selection on tissue specific CREs in the heart, skin, lung, and liver.

Finding a putative selective sweep specific to Europeans led us to ask whether the *ANTXR2* locus differentiates CEU from other human populations. To address this, we calculated relative population differentiation (Fst) between Europeans and either East Asians or Africans. The entire *ANTXR2* locus showed elevated differentiation between European and non-European populations, with a median value of 0.14-0.33 (64th-96th percentile) (**Fig. 4A**). The greatest signal of differentiation occurred outside of the putative selective sweep, but overlapped upstream CREs (including CRE2) that diverged between human and non-human primates (**Fig. 4B**). One particularly interesting haplotype, directly adjacent to the region of high CLR, had a very high Fst in Europeans relative to all other ethnic groups. This region included several SNPs overlapping CREs upstream of *ANTXR2*, including rs41407844—the allele frequency of which was correlated with reported variation in anthrax toxin sensitivity in lymphoblastoid cell lines (*45*): high frequency of the derived allele in Europeans (∼0.85), intermediate in East Asians (∼0.38), and low frequency in African populations (∼0.17; **Fig. 4C-D**). This result, and the observation of elevated Fst at CRE3, CRE4, and CRE9 in the absence of high CLR profile, suggests that Europeans have maintained genetic separation at loci derived from both incomplete soft sweeps on genetic variation from early human divergence and a hard sweep associated with recent European anthrax exposure.

**Figure 4.**
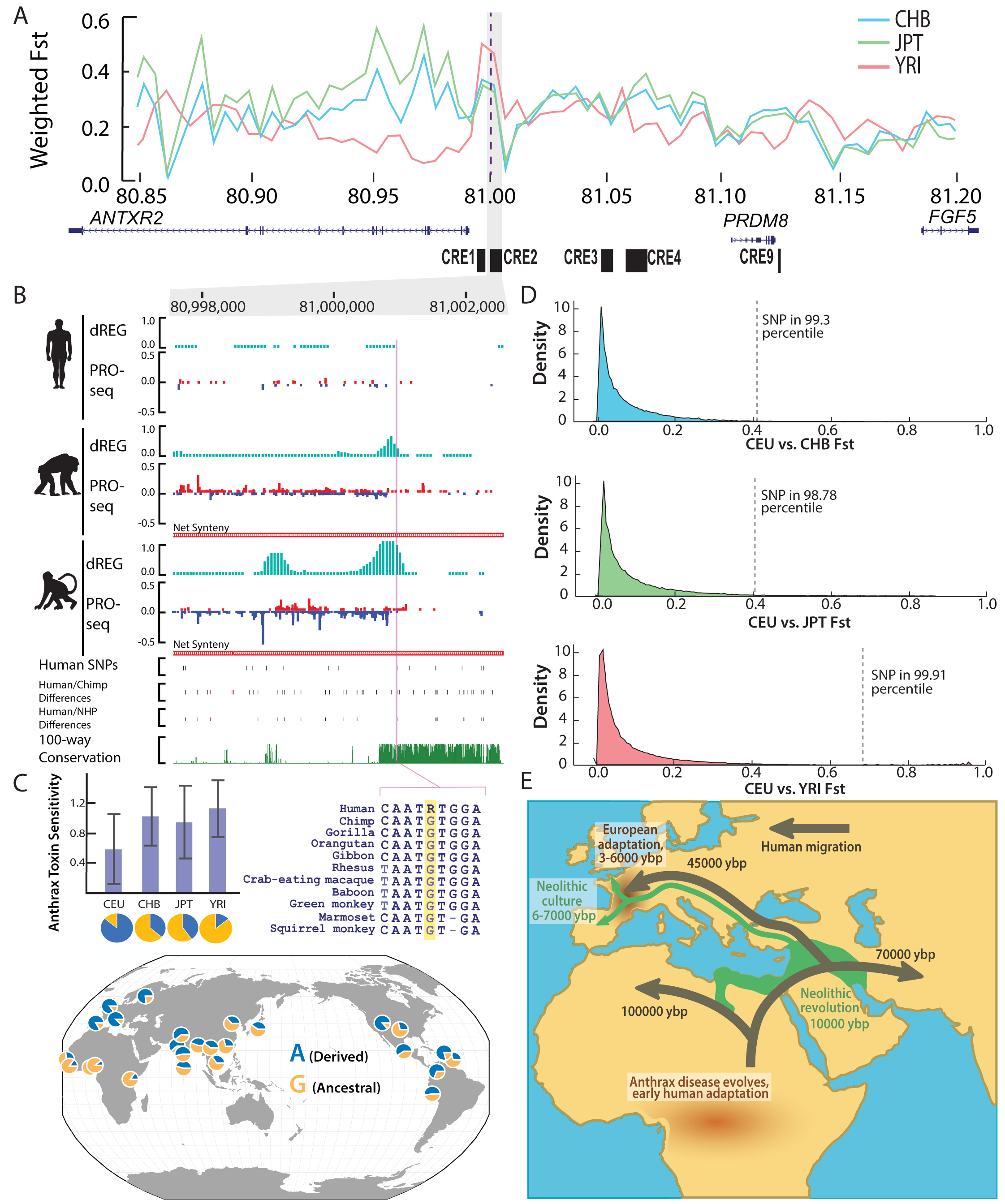
Genetic differentiation in European populations consistent with multiple phases of selection driving changes in *ANTXR2*. **A)** Weighted Fst comparisons between CEU and CHB, JPT, and YRI HapMap populations at the *ANTXR2* locus show a peak around *ANTXR2*. **B)** PRO-seq (normalized by reads per million) and dREG signal at CRE2. rs41407844 falls within a dREG peak shared by chimpanzee and rhesus macaque and the ancestral allele G is conserved within the primate lineage. Conservation of CRE2 between humans and non-human primate tracks show human-specific SNPs, human/chimpanzee differences (SNPs in black, INDELS in red), human/non-human primate differences (SNPs in black, INDELS in red), and 100-way conservation. **C)** Martchenko et al. (45) showed that CEU lymphoblastoid cell lines have the lowest sensitivity to anthrax toxins. rs41407844 allele frequencies across populations. **D)** rs41407844 is above the 98% percentile for Fst comparisons between CEU and CHB, JPT, and YRI HapMap populations. **E)** Model demonstrating the pattern of human migration (dark green) compared to the spread of Neolithic culture (light green) and anthrax disease distribution (brown).

Our work is consistent with the hypothesis that humans adapted to anthrax exposure during several periods of human evolution (**Fig. 4E**). First, ecological changes associated with increased hunting and scavenging early in the evolution of modern humans may have led to increased interactions with *Bacillus* species in Africa. Indeed, *Bacillus* species are believed to have originated in Africa (*55*), and remain endemic to Africa, where they are a source of mortality for wildlife, including wild chimpanzees (*13, 14*). It stands to reason that human ancestors who adopted hunting and scavenging behaviors, especially of ruminant species in endemic areas, would have faced a higher burden of anthrax disease than neighboring chimpanzee relatives. Second, we found evidence of more recent selective pressures acting in Europeans that are consistent with historical records of recent outbreaks within Europe (*18, 43, 56*). The nature of human interactions with anthrax disease might make the *ANTXR2* locus particularly susceptible to soft selective sweeps over time. Both prehistoric and modern anthrax exposure was likely highly sporadic and caste specific—dependent upon a variety of factors including regional endemic areas of the globe, the availability of ruminant species, and the roles of individuals within the population. Indeed, anthrax disease disproportionately affected agricultural and textile workers during the 19th century (*10*). Moreover, *ANTXR2* expression was slightly higher in blood cells isolated from hunter gatherers than in nearby agricultural populations within Africa (*28*), potentially consistent with agricultural populations being at higher risk of exposure to anthrax disease. In summary, our findings imply a complex evolutionary history in the *ANTXR2* locus, supporting a model in which continued hard and soft selection on multiple alleles has continued to drive changes in *ANTXR2* expression within human populations.

## Materials & Methods

### EXPERIMENTAL METHODS

#### Isolation of CD4+ T cells from humans and non-human primates

All human and animal experiments were done in compliance with Cornell University IRB and IACUC guidelines. We obtained peripheral blood samples (60–80 mL) from healthy adult male humans, chimpanzees, and rhesus macaques. Informed consent was obtained from all human subjects. To account for within-species variation in gene transcription we used three individuals to represent each primate species. Blood was collected into purple top EDTA tubes. Human samples were maintained overnight at 4C to mimic shipping non-human primate blood samples. Blood was mixed 50:50 with phosphate buffered saline (PBS). Peripheral blood mononuclear cells (PBMCs) were isolated by centrifugation (750× g) of 35 mL of blood:PBS over 15 mL Ficoll-Paque for 30 minutes at 20C. Cells were washed three times in ice cold PBS. CD4+ T-cells were isolated using CD4 microbeads (Miltenyi Biotech, 130-045-101 [human and chimp], 130-091-102 [rhesus macaque]). Up to 10^8 PBMCs were resuspended in binding buffer (PBS with 0.5% BSA and 2mM EDTA). Cells were bound to CD4 microbeads (20uL of microbeads/107 cells) for 15 minutes at 4C in the dark. Cells were washed with 1–2 mL of PBS/BSA solution, resuspended in 500uL of binding buffer, and passed over a MACS LS column (Miltenyi Biotech, 130-042-401) on a neodymium magnet. The MACS LS column was washed three times with 2mL PBS/BSA solution, before being eluted off the neodymium magnet. Cells were counted in a hemocytometer.

### Luciferase assays

Genomic DNA was isolated from human, chimp, and rhesus macaque PBMCs depleted for CD4+ cells using a Quick-DNA Miniprep Plus Kit (#D4068S; Zymo research) following the manufacturer’s instructions. Putative enhancer regions were amplified from the genomic DNA, restriction digested with KpnI and MluI, and cloned into the pGL3-promoter vector (Promega). The same orthologous regions were amplified from all three species with identical primers where possible or species-specific primers covering orthologous DNA in diverged regions. The media (RPMI-1640) was changed in Jurkat cells one day before transfection. RPMI-1640 with 20% FBS was equilibrated in plates in a 37C incubator prior to transfection. On the day of transfection, Jurkat cells were centrifuged at 100xg for 10 minutes, washed with PBS, and centfrigured again. After centrifugation, Jurkat cells (2 million per reaction) were resuspended in 100 ul room temperature Mirus electroporation solution. Vectors were co-transfected with pRL-SV40 Renilla (Promega) in a 20:1 ratio (2ug pGL3 to 100ng pRL-SV40). Electroporation was done in a Lonza Nucleofector 2b device using program X-001. Immediately after electroporation, 1 ml equilibrated media was added to the cuvette and then the cell mixture was added to 6 well plates and incubated at 37C. 18 hours post-transfection, luminescence was measured in triplicate using the Dual-Luciferase® Reporter Assay System (Promega).

#### CRISPRa in K562

##### Generation of dCas9-KRAB K562 line

Lentivirus was made using lipofectamine 3000 from Invitrogen. Phoenix Hek cells (grown in DMEM with 10% FBS and antibiotics) were seeded in a 6-well plate at 400,000 cells/plate. Cells were grown until ∼90% confluent. 1ug of pHAGE_EF1a_dCas9-KRAB plasmid from addgene (#50919) plasmid was transfected. 24 hours later 3ml/well of virus was mixed with 10ug/ml polybrene and incubated for 5 minutes at room temperature. This mix was added to 300,000 K562 cells and centrifuged for 40 minutes at 800g at 32C. 12–24 hours later the virus was removed and fresh media was added. 24–48 hours later the cells were selected with 150ug/ml Hygromycin B for 2 weeks. The K562 dCas9-KRAB stable cell lines was grown and maintained in Hygromycin B.

##### sgRNA cloning

Primers for sgRNAs were designed using ChopChop (http://chopchop.cbu.uib.no/) for the *ANTXR2* gene and scrambled controls. Primers were located −400 to −50 bp away from the TSS for CRISPRa. A G was added at the 5’ end of primers for use with a U6 promoter, along with restriction sites for cloning. Forward and reverse sgRNAs were synthesized separately by IDT and annealed. T4 Polynucleotide Kinase (NEB) was used to phosphorylate the forward and reverse sgRNA during the annealing. 10× T4 DNA Ligase Buffer, which contains 1mM ATP, was incubated for 30 minutes at 37°C and then at 95C for 5 minutes, decreasing by 5°C every 1 minute until 25°C. Oligos were diluted 1:200 using Molecular grade water. sgRNAs were inserted into the pLenti SpBsmBI sgRNA Hygro plasmid from addgene (#62205) by following the authors protocol (26501517). The plasmid was linearized using BsmBI digestion (NEB) and purified using gel extraction (QIAquick Gel Extraction Kit). The purified linear plasmid was then dephosphorylated using Alkaline Phosphatase Calf Intestinal (CIP) (NEB) to ensure the linear plasmid did not ligate with itself. A second gel extraction was used as before to purify the linearized plasmid. The purified dephosphorylated linear plasmid and phosphorylated annealed oligos were ligated together using the Quick Ligation Kit (NEB). The ligated product was transformed into One Shot Stbl3 Chemically Competent E. coli (ThermoFisher Scientific). 100ul of the transformed bacteria were plated on Ampicillin (200ug/ml) plates. Single colonies were picked, sequenced, and the plasmid was isolated using endo free midi-preps from Omega.

##### Transfection of sgRNA plasmid

The day prior to transfection, the media (RPMI-1640 with 10% media) was changed in K562 cells and cells were diluted to a concentration of 1 million cells/mL. On the day of transfection, K562 cells were centrifuged at 100xg for 10 minutes, washed with PBS, and centrifuged again. RPMI-1640 with 10% FBS was equilibrated in plates in a 37C incubator prior to transfection. After centrifugation, K562 cells (1 million per reaction) were resuspended in 100 ul room temperature Mirus electroporation solution. 2ug of the sgRNA plasmid was added to each reaction. Electroporation was done in a Lonza Nucleofector 2b device using the program for K562 cells. Immediately after electroporation, 1 ml equilibrated media was added to the cuvette and then the cell mixture was added to 6 well plates and incubated at 37C. 12 hours after transfection, 2ug/ml doxycycline was added to cells to activate dCas9-KRAB expression. To confirm overexpression of *ANTXR2*, 4 hours after the addition of doxycycline a portion of the cells were collected for RNA extraction using Trizol. cDNA was generated from RNA samples using the Thermo Fisher High Capacity RNA-to-cDNA kit and qPCR was performed using SsoAdvanced Universal SYBR Green master mix with primers to assay *ANTXR2* expression.

#### Toxin viability assays

K562 cells transfected with CRISPRa plasmids were confirmed to have overexpression of *ANTXR2*. After confirmation and within the 12 hour half-life window of the activating doxycycline, 50,000 K562 cells were added to wells of 96-well plates in 100 ul RPMI-1640, supplemented with 10% RPMI. Anthrax toxin PA (List Biological Laboratories 171D) was added to wells at a concentration of 1ug/ml (or vehicle control). FP59, a recombinant anthrax lethal factor fused to the Pseudomonas Exotoxin A Catalytic Domain (Kerafast ENH013), which is capable of killing blood cells, was added at a concentration of 50ng/ml (or vehicle control). 20 hours after the addition of PA and FP59, 10ul of Alamar Blue (Thermo Fisher #DAL1025) was added to each well. Four hours after the addition of Alamar Blue, fluorescence was measured using a plate reader with an excitation of 570 and emission of 610.

#### Activation of CD4+ T cells

CD4+ T cells were isolated using the above procedure for two human subjects. After equilibration in RPMI-1640 with 10% FBS, cells were stimulated with 25ng/mL PMA and 1mM Ionomycin (P/I or π) or vehicle control (2.5uL EtOH and 1.66uL DMSO in 10mL of culture media). Thirty minutes after activation or addition of the vehicle control, cells were treated with 2.5ug/ml Recombinant PA (List Biological Laboratories 171D) and 500ng/ml Recombinant LF (List Biological Laboratories 172A) or vehicle control. 24 hours later, media from non-activated, non-activated with toxin treatment, activated, and activated with toxin treatment wells was collected for ELISA.

#### IL-2 ELISA on CD4+ T cells

ELISA was done using the R&D Human IL-2 DuoSet Kit (Catalog #DY202).

Plate preparation—16 hours prior to the ELISA experiment, the capture antibody was added to 96-well plates at a concentration of 0.5ug/ul in 100ul of PBS. Plates were sealed and left at room temperature. The following day, the diluted capture antibody was aspirated and washed with 400ul Wash Buffer three times. 300ul of Block Buffer was added to each well and incubated for at least one hour. After incubation, plates were washed with 400ul Wash Buffer.

ELISA assay—100ul of RPMI from the samples or standards diluted in the Reagent Diluent were added to the prepared 96-well plate. The plate was covered and left to incubate for 2 hours at room temperature. 100ul of the Detection Antibody diluted at 1:60 with the Reagent Diluent was added to each well. The plate was washed with Wash Buffer. 100ul of Streptavidin HRP diluted at 1:40 in the Reagent Diluent was added to each well. The plate was covered, protected from light, and incubated for 20 minutes at room temperature. The plate was washed and 100ul of the Substrate Solution was added to each well, the plate was covered, protected from light, and incubated for 20 minutes at room temperature. 50ul of Stop Solution was added to each well and mixed. Fluorescence was measured on a plate reader at 450nm and wavelength corrected at 540nm.

#### Hi-C library preparation

##### Cell preparation

CD4+ T cells were isolated according to the above procedure. After >1 hour of equilibration in RPMI-1640 supplemented with 10% FBS, cells were centrifuged at 300xg, washed with PBS, and centrifuged again. Cells were resuspended in a mixture of 1% paraformaldehyde in 1x PBS. Cells were incubated at room temperature for 10 min on a rocker. Paraformaldehyde was quenched by the addition of 2.5M Glycine to a Cf=0.2M. Cells were incubated for room temperature for 5 minutes on a rocker. Cells were centrifuged at 4C, washed in cold PBS, centrifuged, and PBS was aspirated. Pellets were flash frozen using dry ice and stored at −80C prior to library preparation.

##### Hi-C

The protocol detailed in (*35*) was followed with the following adjustments. After the addition of lysis buffer, cells were incubated on ice for 30 min. MboI (NEB #R0147) was used for restriction digestion. Following DNA purification, unligated biotin was removed using a mixture of 0.5uL of 10mM dATP, 0.5uL dGTP, 20uL 3000U/ml T4 DNA polymerase (NEB #M02030) for each sample. Samples were incubated for 4 hours at room temperature and then T4 DNA polymerase was inactivated at 72C for 20min. Shearing was done using a Bioruptor sonicator using the LOW setting 30S ON/ 90S OFF for 2 cycles of 10 minutes. Libraries were prepared with the NEBNext Ultra II Library Preparation Kit (NEB #E7103). Samples were sequenced on a combination of Illumina’s NovaSeq 6000 and HiSeq 4000 at Novogene.

## DATA ANALYSIS

### Hi-C analysis and visualization

Individual Hi-C samples were mapped to hg19 (for human samples) or rheMac8 (for rhesus macaque samples) using Juicer (*57*). Replicates for each species were combined using Juicer’s mega function. The combined rhesus macaque aligned dataset was lifted over to hg19 using Crossmap (*58*).

### Mapping orthologs between species

Cross-species comparison of genomic coordinates and genes was based on the methods using in (*34*). Briefly, all datasets for chimpanzee and rhesus macaque were converted to the human assembly (hg19) using CrossMap (*58*). Reciprocal-best (rbest) nets were used to convert genomic coordinates between genome assemblies using (*59*).

### PRO-seq and RNA-seq differential expression

PRO-seq data from human, chimpanzee, and rhesus macaque were published in ref (*34*). We mapped PRO-seq reads using standard informatics tools. Our PRO-seq mapping pipeline begins by removing reads that fail Illumina quality filters and trimming adapters using cutadapt with a 10% error rate. Reads were mapped with BWA (*60*) to the appropriate reference genome (either hg19, panTro4, or rheMac3) and a single copy of the Pol I ribosomal RNA transcription unit (GenBank ID# U13369.1). Mapped reads were converted to bigWig format for analysis using BedTools (*61*) and the bedGraphToBigWig program in the Kent Source software package (*62*). The location of the RNA polymerase active site was represented by the single base, the 3′ end of the nascent RNA, which is the position on the 5′ end of each sequenced read.

### ANTXR2 RNA-seq analysis in PBMCs

Bakiga and Batwa PBMC RNA-seq data (*28, 29*)) was downloaded for the control condition (GEO series GSE120502). Rhesus macaque RNA-seq data was downloaded for the control condition (NCBI BioProject PRJNA246101). RNA-seq data was mapped using Salmon (*63*) and NCBI RefSeq genes to hg19. Rhesus macaque RNA-seq data was mapped to hg19 using the same parameters. *ANTXR2* transcripts per million were compared between samples for transcript NM_001145794.1.

### DICE RNA-seq and eQTL analysis

RNA-seq from 85 human CD4+ naive T cell samples from DICE (*27*) was downloaded under dbGap protocol 23187. RNA-seq data was mapped using Salmon (*63*) and NCBI RefSeq genes to hg19. Rhesus macaque RNA-seq data was mapped to hg19 using the same parameters. *ANTXR2* transcripts per million were compared between samples for transcript NM_001145794.1. We retrieved eQTLs for *ANTXR2* from the DICE online database.

### Sweepfinder2 CLR scan

Human genome data was taken from phase3 of the 1,000 Genomes Project(*64*). VCFs were subsetted based on their population group. The b-value maps that are used to compute the effect of background selection on the human genome were taken from(*65*). Recombination maps from the deCODE database(*66*) were used. Sweepfinder2(*46*) was used to calculate the composite-likelihood-ratio for CEU, JPT, CHB, and YRI on chromosome 4.

### Fst analysis

All Fst values were computed using VCFtools (*67*) using the --weir-fst command with a window size of 5kb and a step size of 5kb. In addition to the Fst calculations for bins of 5kb, Fst was calculated for each SNP of chromosome 4. Fst was calculated for all pairwise comparisons of CEU, YRI, JPT and CHB.

## Supporting information

Merged Supplement

## Acknowledgements

We thank A. Siepel, H. Hijazi, A. Clark, B. Marks, and all members of the Danko lab for valuable discussions and suggestions. New Hi-C and RNA-seq data from human and rhesus macaque CD4+ T-cells have been deposited in GEO, and an accession number will be added shortly. Work in this publication was supported by R01-HG010346 and R01-HG009309 (NHGRI) to CGD and an F31-1AI140050 (NIAID) to LAC. The content is solely the responsibility of the authors and does not necessarily represent the official views of the US National Institutes of Health.

