## Supplementary material for "Multiple stages of evolutionary change in anthrax toxin receptor expression in humans": Merged Supplement

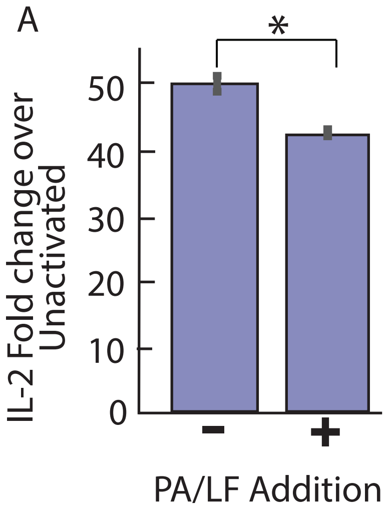


**Supplementary Figure 1. Effect of anthrax toxin treatment on T cell activation. A)** CD4+ T cells produce less IL-2 upon activation with PMA and ionomycin when treated with anthrax toxins protective antigen (PA) and LF as measured by ELISA.

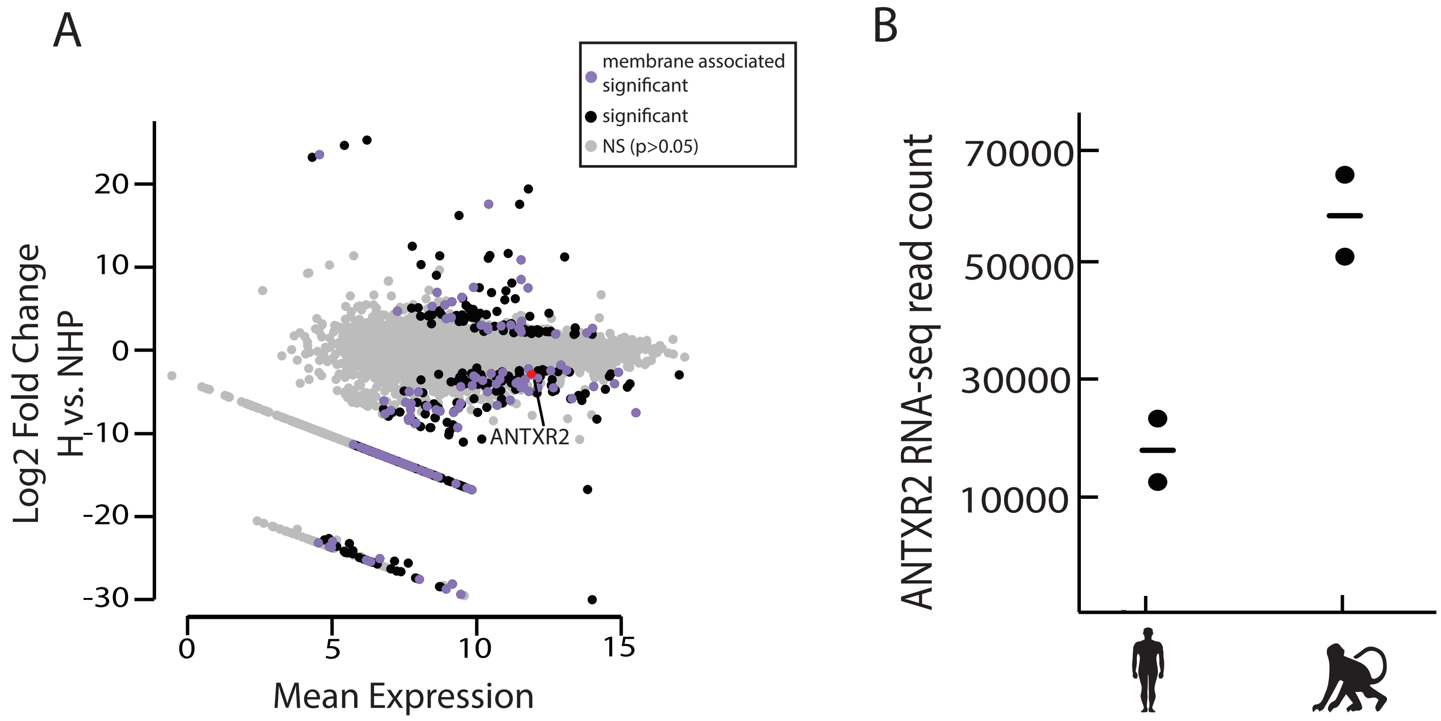


**Supplementary Figure 2. Differential RNA expression between humans and non-human primates. A)** RNA-seq comparison of protein-coding genes between humans and non-human primates in CD4+ T cells. The GO term ‘integral component of the membrane’ is enriched in differentially transcribed genes. **B)** *ANTXR2* RNA-seq in CD4+ T cells is consistent with the relative levels of *ANTXR2* transcription based on PRO-seq with higher expression in rhesus macaque.

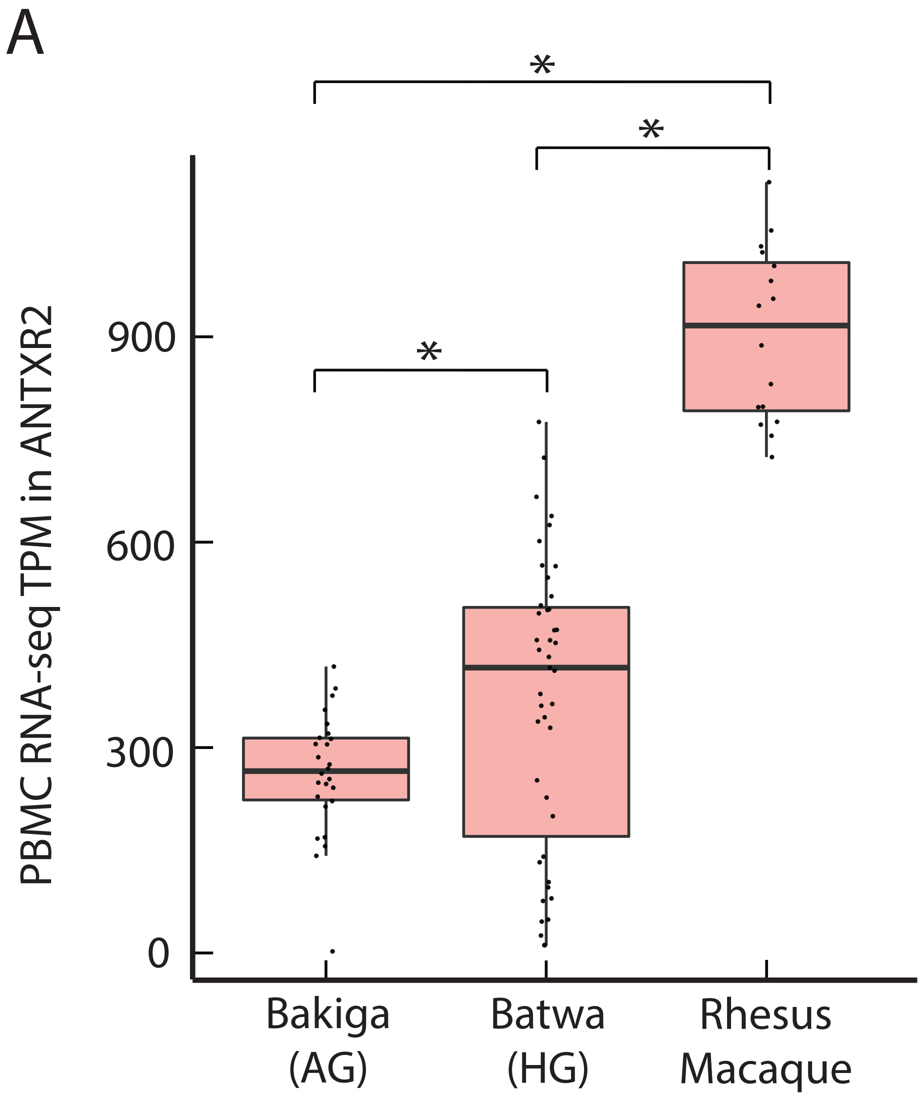


**Supplementary Figure 3. Comparison of ANTXR2 RNA expression in PBMCs. A)** RNA-seq in PBMCs from Bakiga (agricultural) and Batwa (hunter-gatherer) individuals compared to rhesus macaque PBMCs. *ANTXR2* expression is significantly different for each comparison based on a Wilcoxon rank sum test (Bakiga vs. Batwa p=0.01; Bakiga vs. rhesus macaque p=1.2x10^-11; Batwa vs. rhesus macaque p=1.3x10^-13).

A

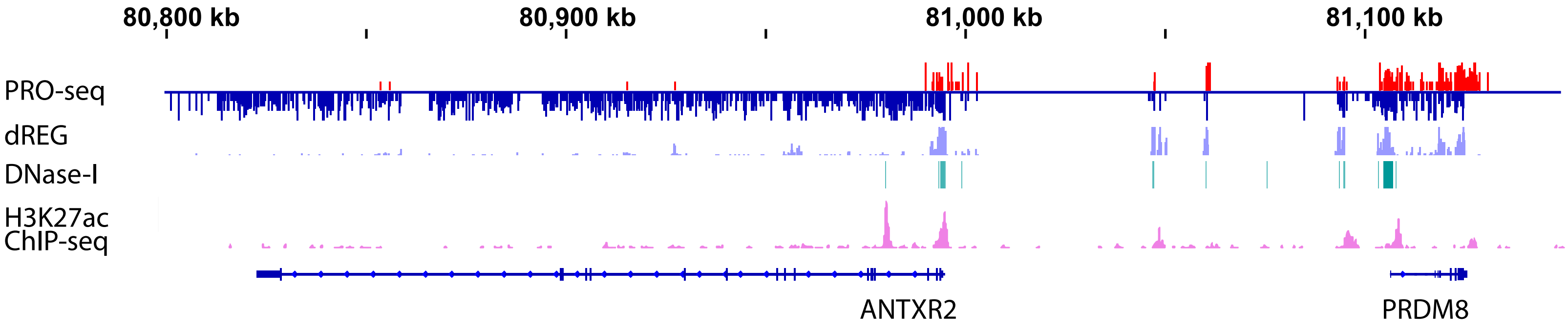


**Supplementary Figure 4. H3K27ac and DNase-I-seq at *ANTXR2*. A)** PRO-seq, dREG signal, DNase-I-seq peaks, and H3K27ac ChIP-seq at the *ANTXR2* locus.

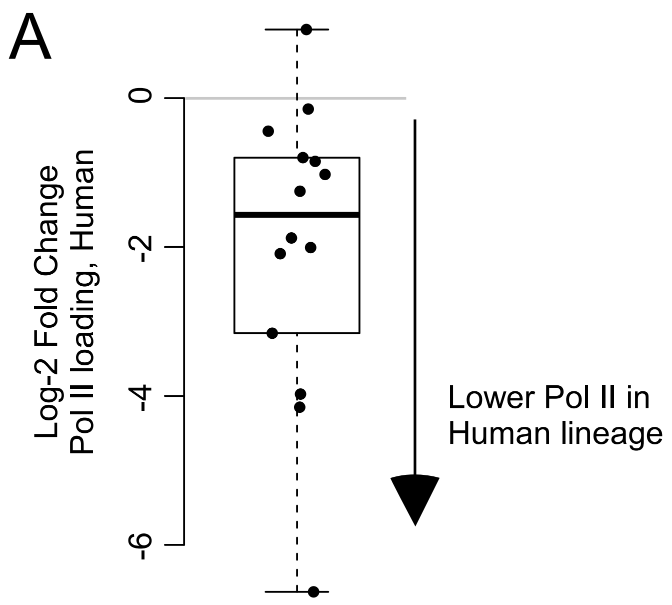


**Supplementary Figure 5. Decrease Pol II Loading on *ANTXR2* CREs. A)** Fourteen CREs in the *ANRXR2* locus show a bias for decreased Pol II loading in humans compared with non-human primates (chimpanzee and rhesus macaque). The Y axis denotes the log-2 fold-change in human. Values below 0 indicate decreased expression.

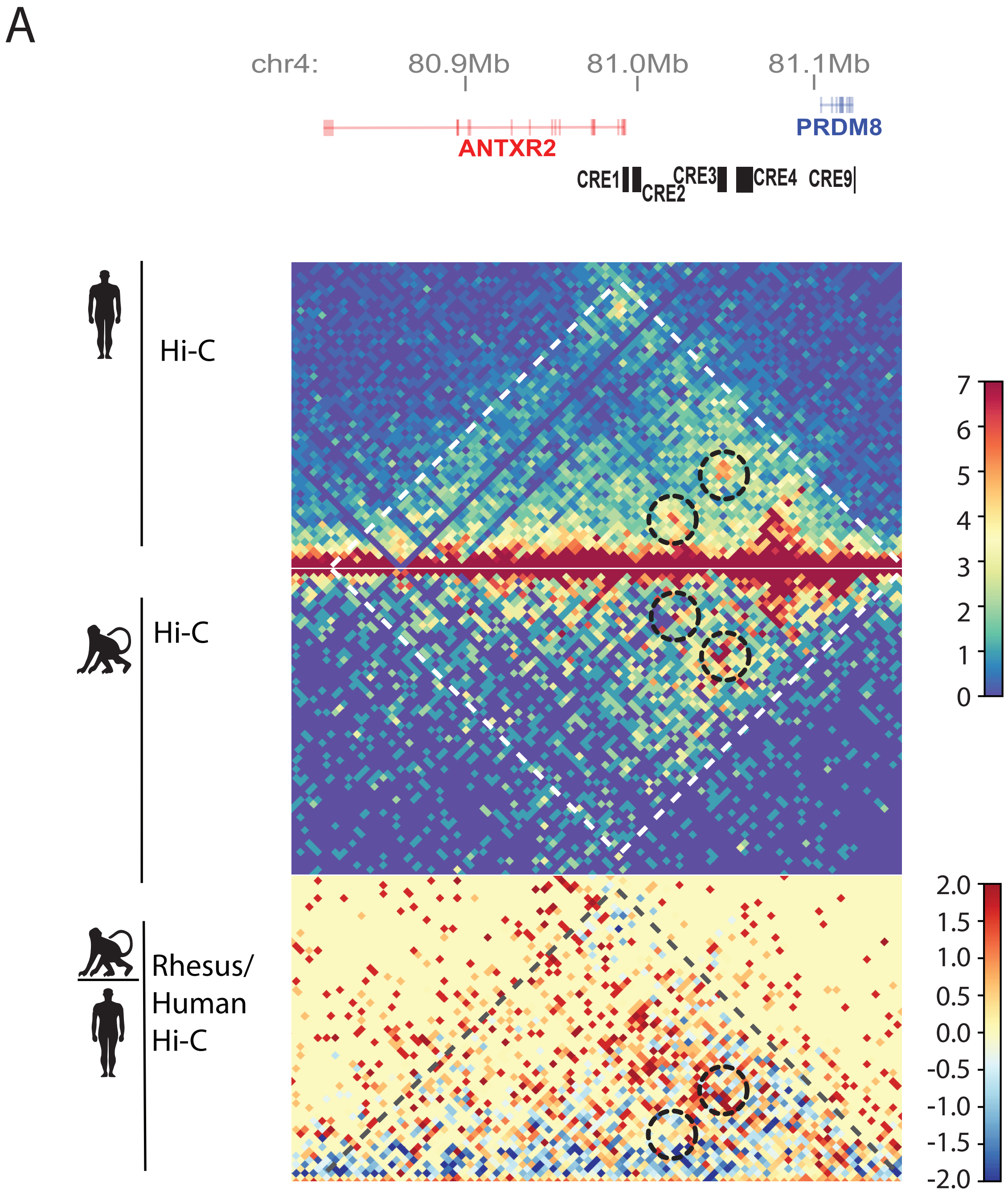


**Supplementary Figure 6. Comparison of human and rhesus macaque Hi-C. A)** Rhesus macaque Hi-C signal divided by human Hi-C shows an increase in contacts within *ANTXR2*'s TAD. TADs are marked with white dotted lines (grey for human/rhesus macaque). Focal contacts with the *ANTXR2* promoter are circled with black dotted lines.

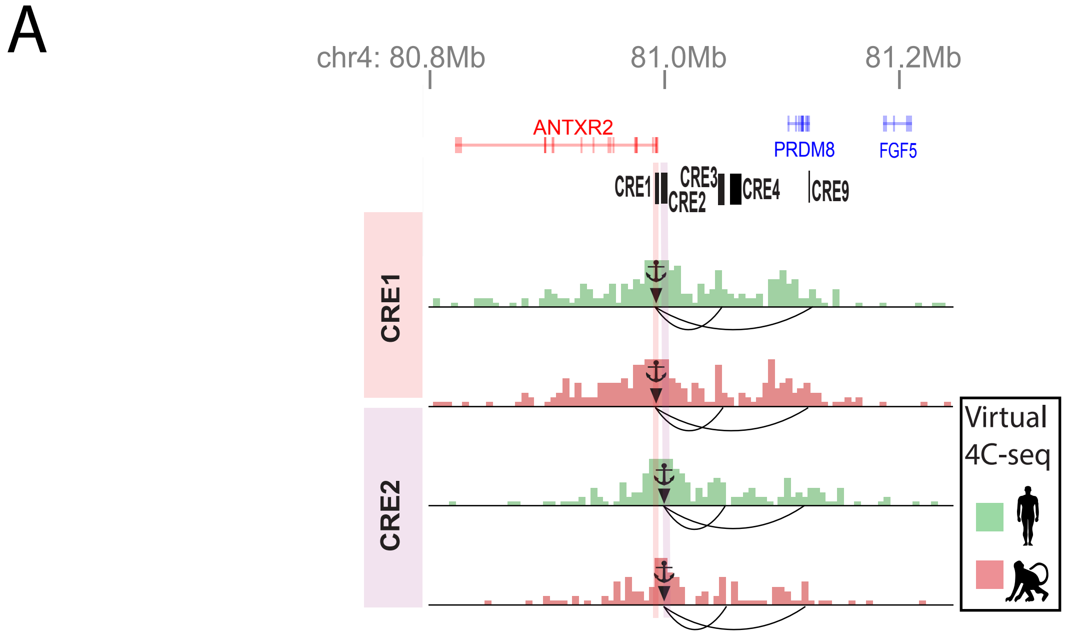


**Supplementary Figure 7. Virtual 4C-seq of proximal CREs. A)** Virtual 4C-seq plots of CRE1 and CRE2 show contact with the upstream CREs that had significantly higher activity in non-human primates.

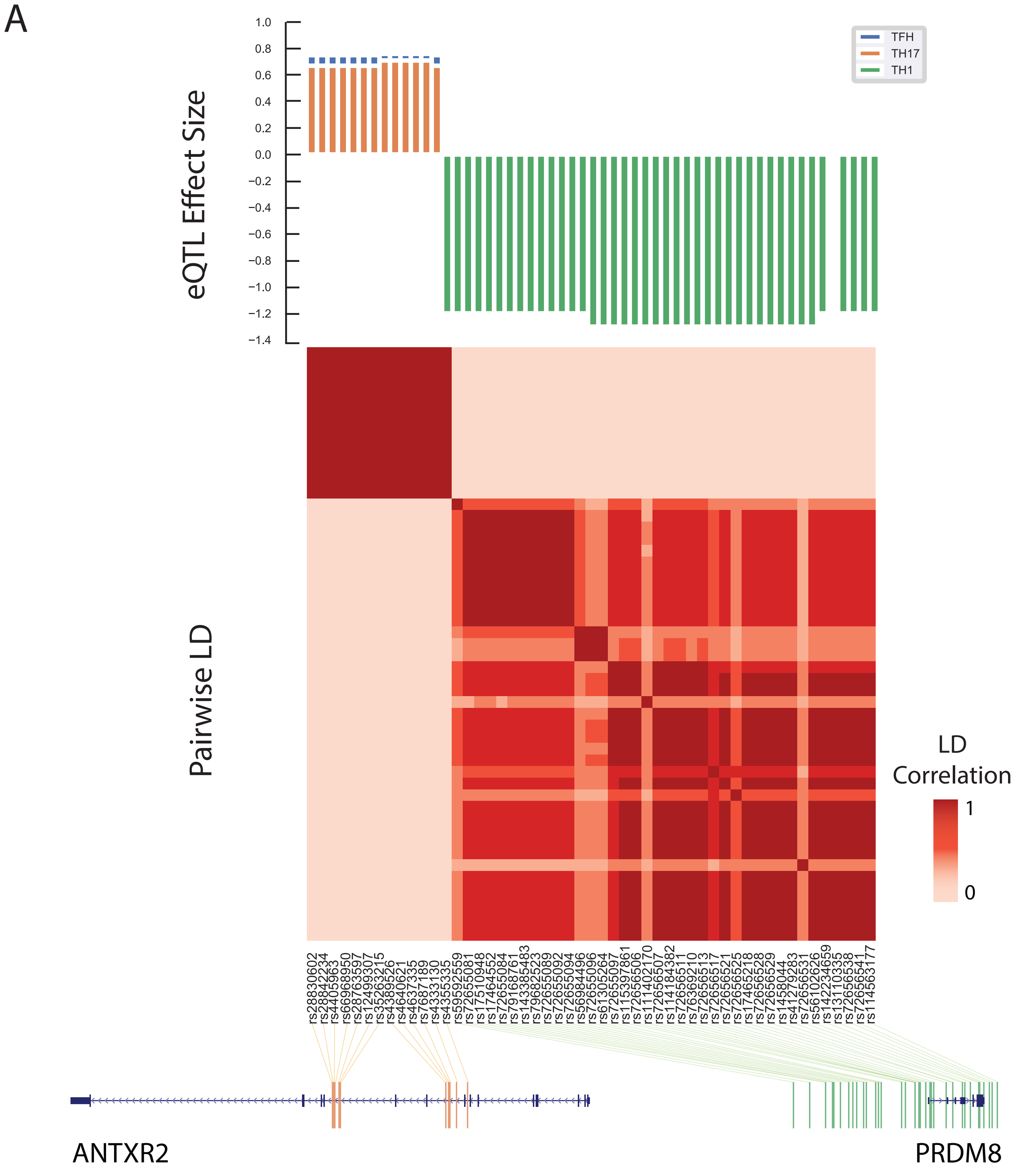


**Supplementary Figure 8. *ANTXR2* eQTLs in humans. A)** Expression eQTLs for ANTXR2 fall into two main regions: within the gene and upstream of *ANTXR2* around *PRDM8*. Genic eQTLs are in linkage disequilibrium (LD) and have a positive effect on *ANTXR2* expression. Upstream eQTLs fall within two blocks of LD and have a negative effect on ANTXR2 expression.

A

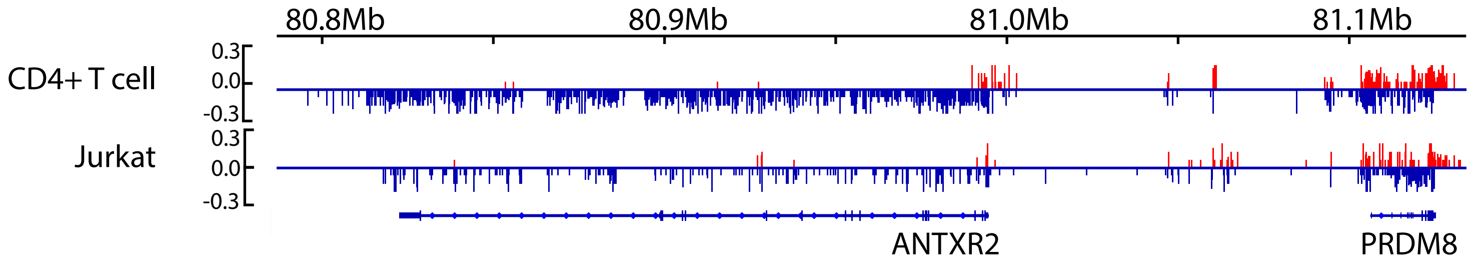


**Supplementary Figure 9. Jurkat vs. CD4 PRO-seq. A)** PRO-seq from Jurkat and human CD4+ T cells shows a similar regulatory landscape around *ANTXR2*.

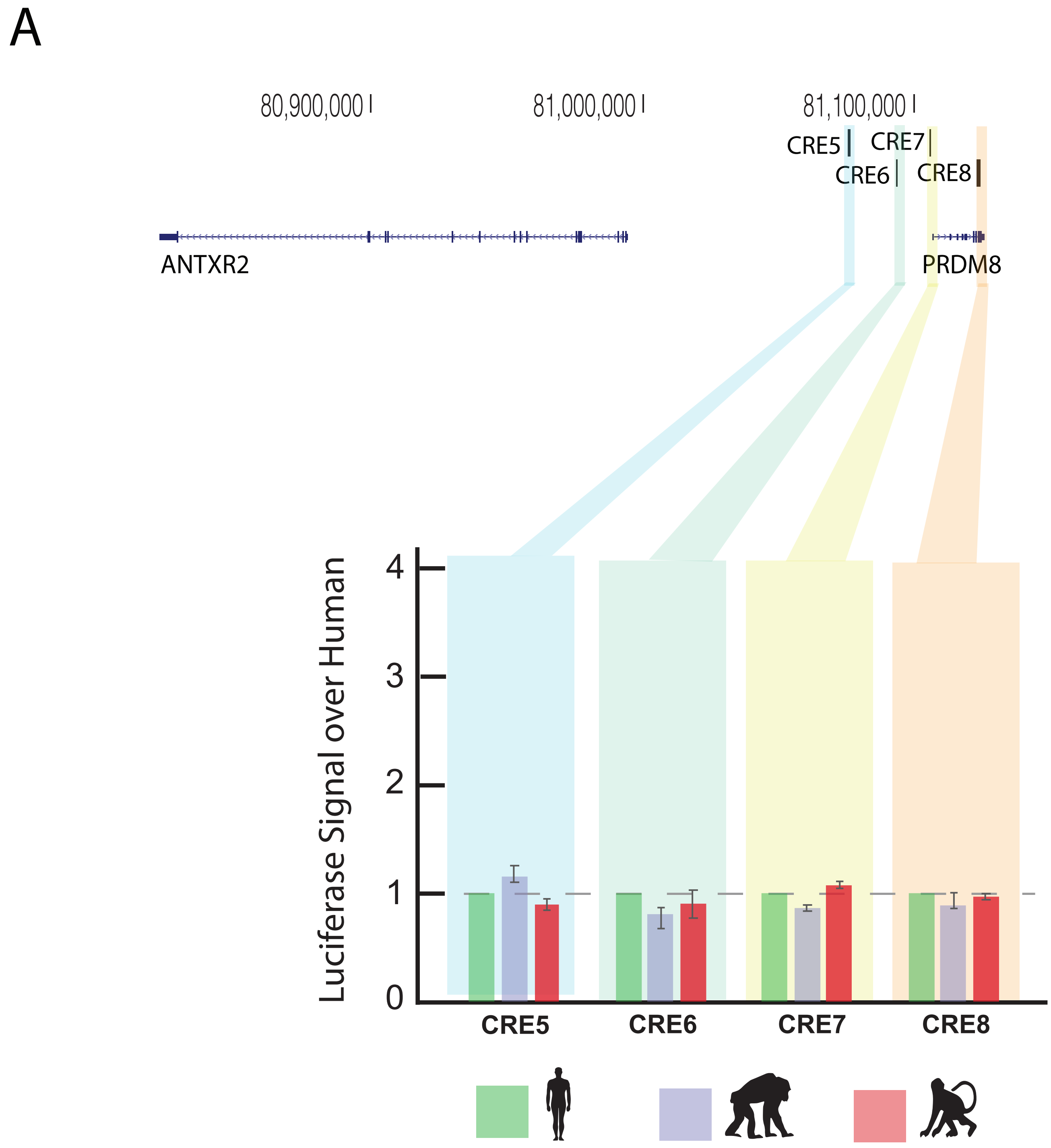


**Supplementary Figure 10. Luciferase data of other CREs. A)** Luciferase assay performed in Jurkat cells to test the activity of regulatory elements in human, chimpanzee, and rhesus macaque shows no significant changes in activity for CREs 5, 6, 7, and 8.

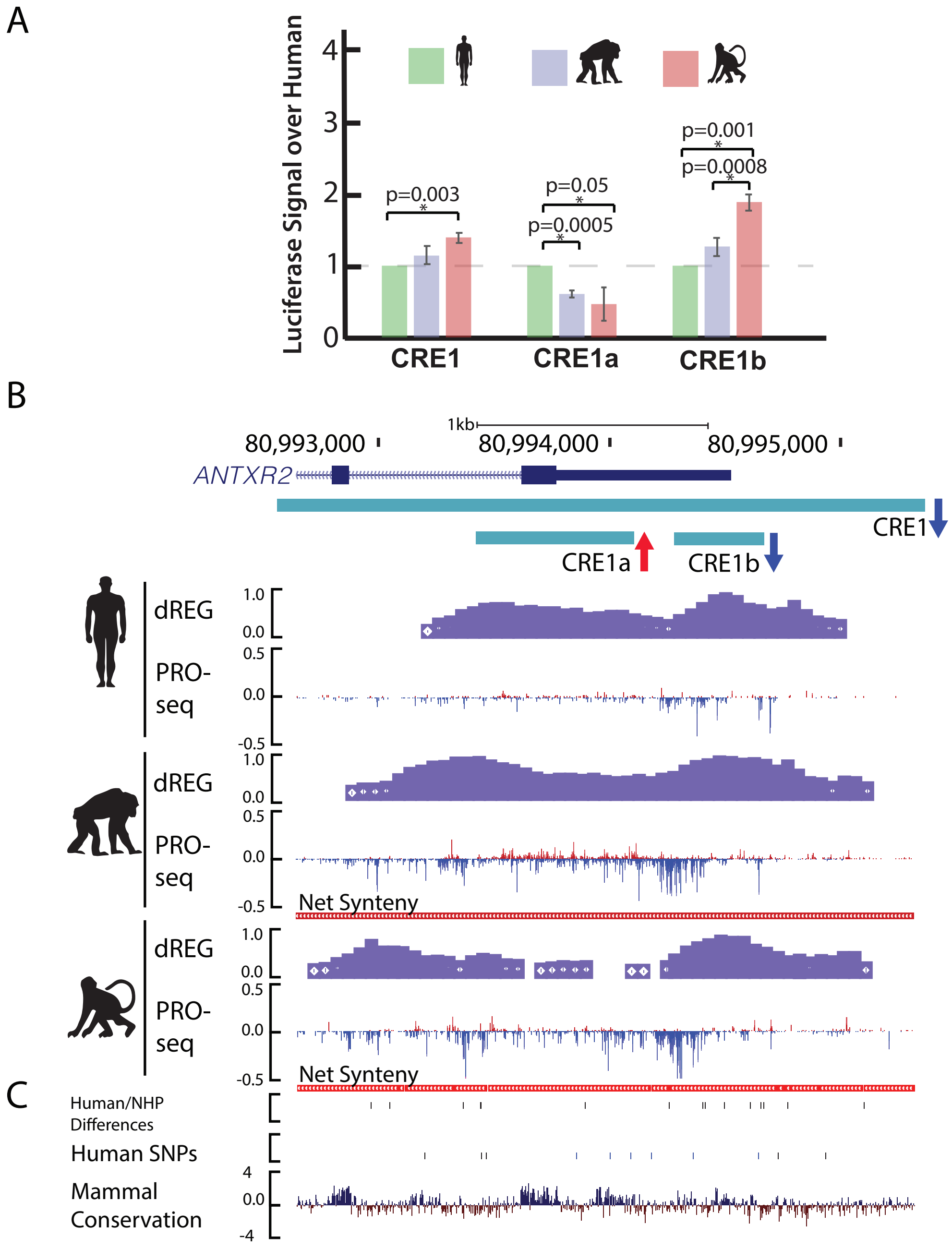


**Supplementary Figure 11. Complex promoter of *ANTXR2*. A)** Luciferase assay performed in Jurkat cells to test the activity of regulatory elements in human, chimpanzee, and rhesus macaque shows a human-specific decrease in the activity of CRE1. Sub-CREs 1a and 1b show a human-specific increase and decrease in activity, respectively. **B)** PRO-seq and dREG signal from human, chimpanzee, and rhesus macaque at CRE1 in the *ANTXR2* promoter. **C)** Human-specific changes, human common SNPs, and PhyloP conservation of the *ANTXR2* promoter.

A

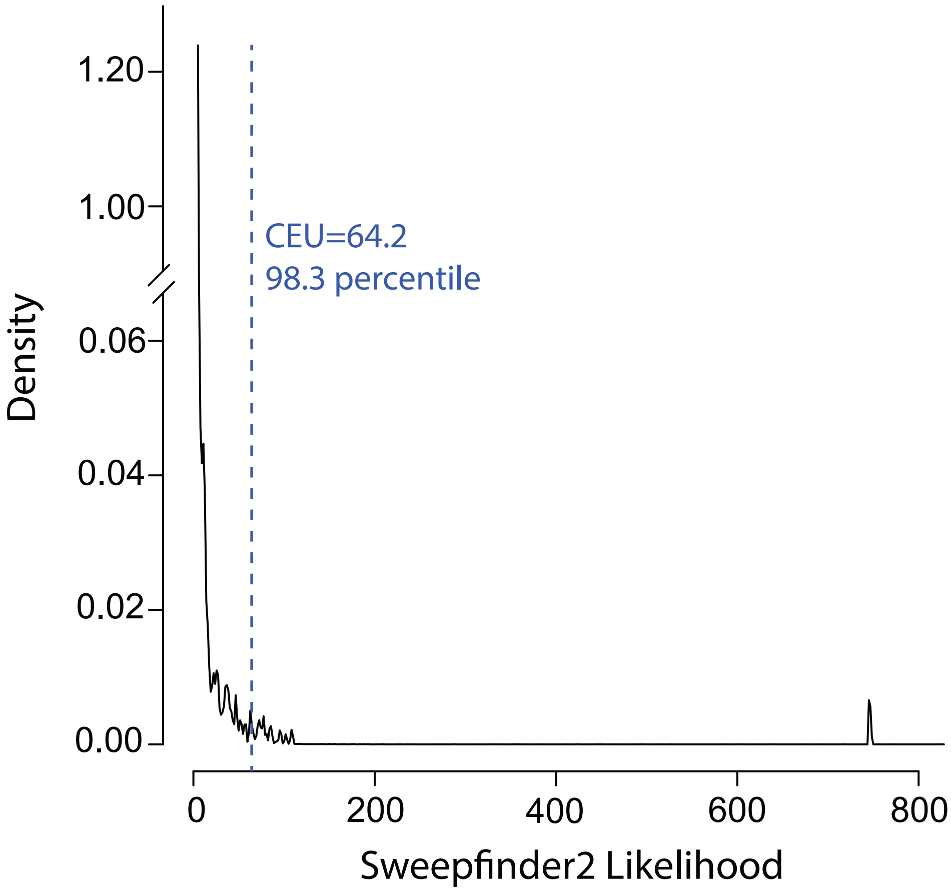


**Supplementary Figure 12. CLR percentile in CEU. A)** The predicted selective sweep upstream of *ANTXR2* falls within the 98th percentile for all Sweepfinder2 likelihoods on chromosome 4.

A

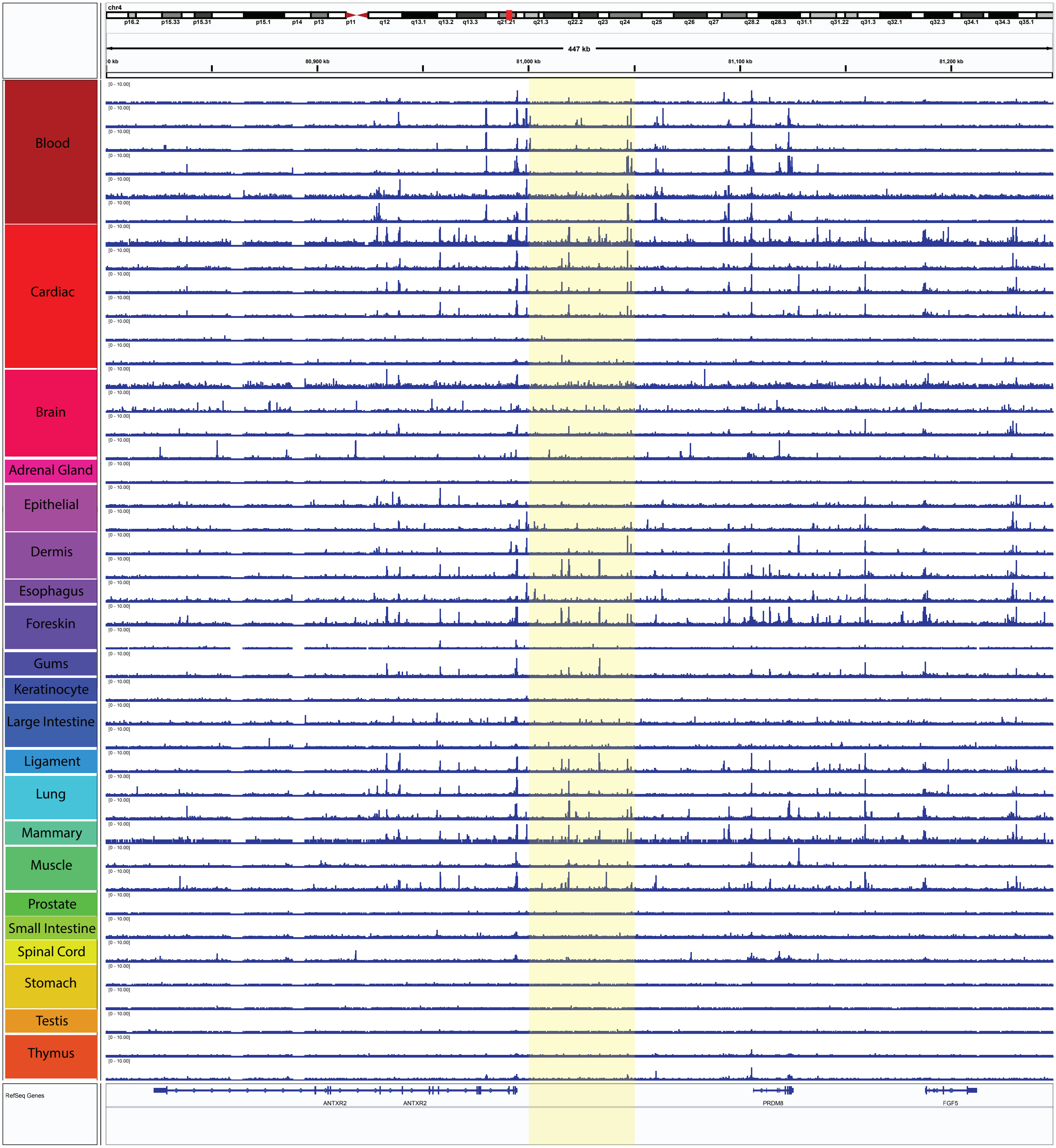


**Supplementary Figure 13. DNase-I-seq profiles across diverse ENCODE tissues at *ANTXR2*. A)** Patterns in DNase-I-seq data reveal differential regulatory landscapes between tissues at the *ANTXR2* locus.
